# Differential requirements of tubulin genes in mammalian forebrain development

**DOI:** 10.1101/304196

**Authors:** Elizabeth Bittermann, Ryan P. Liegel, Chelsea Menke, Andrew Timms, David R. Beier, Beth Kline-Fath, Howard M. Saal, Rolf W. Stottmann

## Abstract

Tubulin genes encode a series of homologous proteins used to construct microtubules which are essential for multiple cellular processes. Neural development is particularly reliant on functional microtubule structures. Tubulin genes comprise a large family of genes with very high sequence similarity between multiple family members. Human genetics has demonstrated that a large spectrum of cortical malformations results from *de novo* heterozygous mutations in tubulin genes. However, the absolute requirement for most of these genes in development and disease has not been previously tested in genetic, loss of function models. Here we present two novel pathogenic tubulin alleles: a human *TUBA1A* missense variant with a phenotype more severe than most tubulinopathies and a mouse ENU allele of *Tuba1a*. Furthermore, we directly test the requirement for *Tuba1a, Tuba8, Tubb2a* and *Tubb2b* in the mouse by deleting each gene individually using CRISPR-Cas9 genome editing. We show that loss of *Tuba8, Tubb2a* or *Tubb2b* does not lead to cortical malformation phenotypes or impair survival. In contrast, loss of *Tuba1a* is perinatal lethal and leads to significant forebrain dysmorphology. Thus, despite their functional similarity, the requirements for each of the tubulin genes and levels of functional redundancy are quite different throughout the gene family. The ability of the mouse to survive in the absence of some tubulin genes known to cause disease in humans suggests future intervention strategies for these devastating tubulinopathy diseases.

## INTRODUCTION

Tubulin proteins are fundamental building blocks of the cell and assemble into dynamic microtubules. Microtubules are especially crucial for cortical development where they are used in multiple cellular contexts such as the mitotic spindle, axons, dendrites, and cilia (1). Mutations in tubulin genes are now known to cause multiple human cortical malformations including lissencephaly, polymicrogyria, microcephaly, and dysmorphic basal ganglia (2). These are collectively discussed as “tubulinopathies.” Many variants leading to malformations of cortical development have now been identified in *TUBULIN, ALPHA-1A (TUBA1A)* (2–18), *TUBULIN, BETA-2A (TUBB2A)* (19–22), *TUBB2B* (4, 5, 15, 23–29), *TUBB4A* (30), and *TUBB/TUBB5* (31). In fact, for *TUBA1A* and *TUBB2B* alone, there are now 71 identified variants in cortical malformation patients (7–23, 27–34). Mutations of *TUBA1A* and *TUBB2B* account for ~5% and ~1.2% of lissencephaly and related malformations of cortical development (32). Both, but especially *TUBA1A*, are associated with a wide spectrum of severity (32). With the exception of one variant inherited from a mosaic parent (6), all of these variants are heterozygous, *de novo* changes in the identified proband and not recessively inherited.

Previous work has shown that both α- and β- tubulins can be separated into different classes (isotypes) and some general characteristics about their spatiotemporal expression domains have been established (33, 34). However, the completed genome sequences of many species including mouse and human have revealed the presence of more tubulin proteins than those indicated by the initial six classes. The requirements and roles in cortical development for some of the individual tubulin genes are still unknown. One possible mechanism of disease in the tubulinopathies is that the missense variants produce monomers which alter normal tubulin function(s) in any number of cellular contexts. This perturbed function could be in assembly dynamics of the microtubule, interaction of the microtubule with the wide array of microtubule associated proteins, perturbation of post-translation modifications of tubulins, disrupting processive movement of motor proteins along the microtubule, and/or formation of microtubule based primary and motile cilia in the brain.

Most tubulin genes exhibit high sequence homology and many are clustered in the genome. Complete genome assemblies now highlight that *TUBB2A* and *TUBB2B* are immediately adjacent to each other in both the human and mouse genome and are virtually identical (443 identical amino acids of 445 total). Similarly, *TUBA1A, TUBA1B*, and *TUBA1C* genes are genomically adjacent and contain high sequence homology in both human and mouse. These characteristics suggest these genes may be the product of genome duplications and the functions of each individual gene may be shared across the cluster(s).

Despite the central importance of these genes in the cytoskeleton and their relevance for human cortical malformation, relatively few genetic models of tubulins exist. No null alleles are published for *Tuba1a, Tuba1b, Tuba1c, Tubb2a*, or *Tubb2b*. ENU mutagenesis efforts have identified alleles in *Tuba1a* (7, 35, 36) and *Tubb2b* (37) and a mouse model of CFEOM for *Tubb3* has been made (38). Here we identify two new alleles of *TUBA1A* in human and mouse. We also test the requirements for *Tuba1a, Tubb2a*, and *Tubb2b* with specific genetic deletions.

## RESULTS

### A novel pathogenic allele of *TUBA1A*

As part of our initiative to identify the genetic causes of congenital malformations of the brain, we were referred an unborn patient with multiple anomalies for genetic testing. The male patient born at 37 weeks to a 32 year old woman and her 34 year old unrelated husband. The pregnancy was complicated by identification of multiple anomalies noted on high-resolution ultrasound, including hydrocephalus, abnormal brainstem, hypoplastic cerebellum with absent vermis, and a large interhemispheric cyst with abnormal sulcation. There were also bilateral clubfeet. A fetal MRI scan performed at 22 weeks confirmed the ultrasound findings and identified several additional findings including aqueduct stenosis and macrocephaly. Non-invasive prenatal testing with free fetal DNA showed that there was no aneuploidy and a fetal echocardiogram was normal. The infant was delivered via Cesarean section, comfort care was provided, and the infant expired at about 8 hours of age.

Examination of the infant post mortem showed a normal sized fetus with enlarged head. There was frontal bossing with a high and wide forehead, the eyes were deeply set and there was a depressed nasal bridge, square face, and mild micrognathia. The nose was short, the philtrum was formed but short, and the mouth was small with thin upper and posterior vermilions. The cheeks were full and the palate was intact. The ears were normally placed and there was overfolding of the superior helix. Chest was symmetric. There was campodactyly of the fingers with no other digital anomalies. Lower extremities were mildly bowed, there was bilateral talipes equinovarus with metatarsus adductus. No additional skeletal anomalies were noted. Genitalia were normal male.

A post-mortem MRI revealed severe lateral ventriculomegaly with bilateral ventricular rupture and absence of the septum pellucidum (Fig. 1A). There was moderate third ventriculomegaly with absence of cerebrospinal fluid (CSF) in the aqueduct of Slyvius in association with a thickened tectum, consistent with aqueduct obstruction (Fig. 1B). There was no evidence of heterotopias along the ventricular surface, although cysts were noted at the caudothalamic ganglionic eminences (Fig. 1A). The supratentorial brain mantle was reduced in size and there was an overall significant loss in white matter volume. The sulcation was shallow without evidence for cobblestone lissencephaly (Fig. 1A). The corpus callosum was nearly absent with thin remnant in the area of the genu. The hippocampal tissue was thin and the thalamus appeared partially fused. The basal ganglia structures were poorly defined with lack of visualization of the internal capsule (Fig. 1C). The brainstem was small with a “Z” shaped configuration (Fig. 1B) and small central cleft. The cerebellar hemispheres and vermis were small with limited sulcation. The optic nerves were hypoplastic and olfactorary bulbs absent. Outside the central nervous system (CNS), we observed mild micrognathia but no evidence of cleft lip/palate.

**Figure 1.**
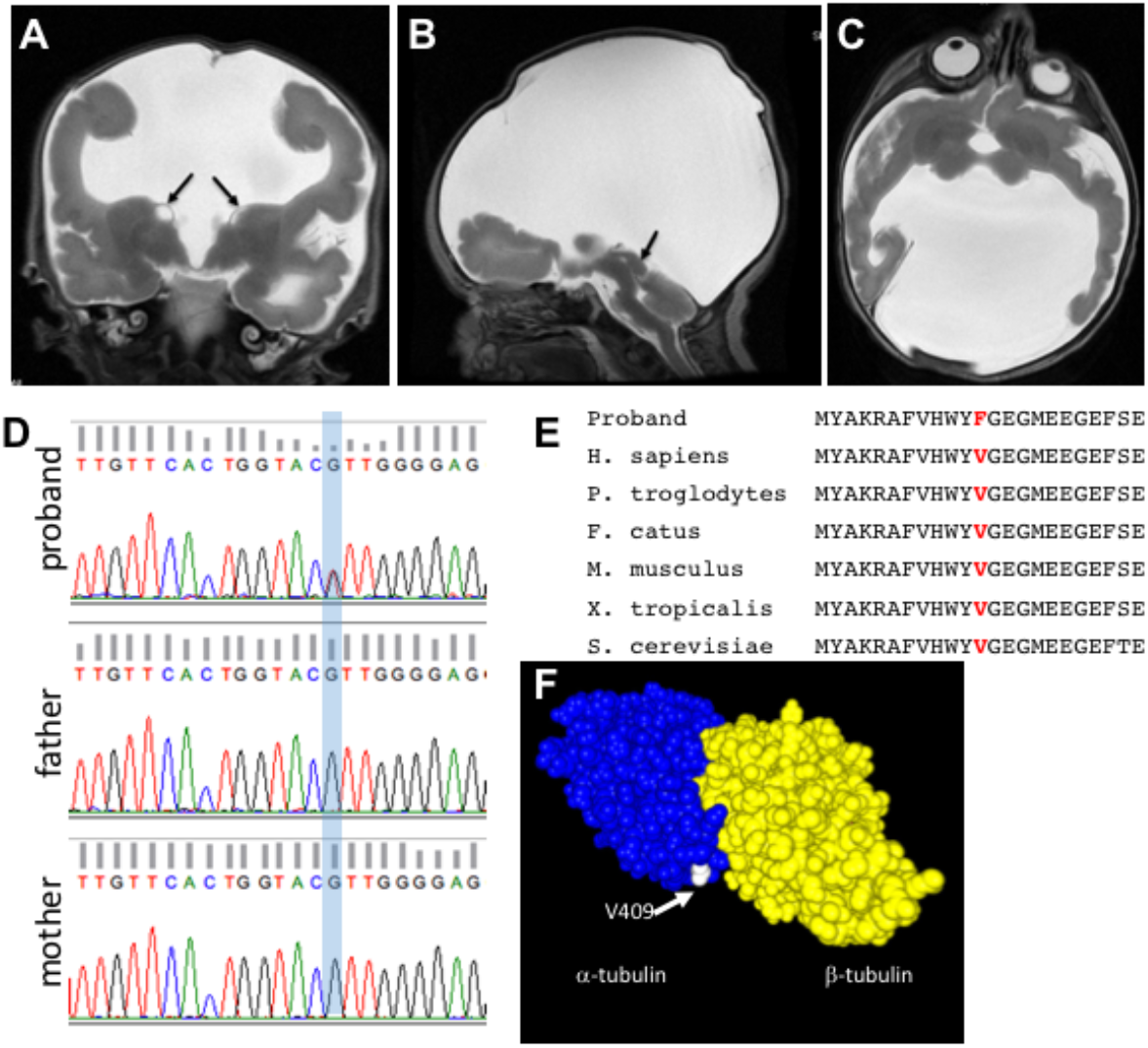
A novel variant in *TUBA1A*. (A-C) MRI Imaging. (A) Coronal T2-weighted MRI at 38 weeks of gestation showing severe lateral ventriculomegaly and moderate third ventriculomegaly with absence of the septum pellucidum. The brain mantle is thin with shallow sulcation. Small cysts are present at the ganglionic eminence present at the caudothalamic groove (arrows). (B) Sagittal T2 image demonstrates lack of CSF distention of the aqueduct of Slyvius with thickened tectum (arrow). The brainstem is small and demonstrates a “Z” shaped configuration. (C) The basal ganglia structures are small with lack of visualization of normal internal capsule. (D) Sanger sequencing validation of the exome sequencing results. Blue shading highlights the heterozygous *TUBAlA* variant. (E) Protein alignment shows the conserved nature of the amino acid residue altered in the p.V409F patient. (F) Three-dimensional modeling of the variant highlights the exterior position of the V409 amino acid (white) on the a-tubulin protein (blue).

### Exome sequencing analysis

Chromosome analysis for the infant showed that he had a maternally inherited balanced translocation, 46,XY,t(11;22)(q23;q11.2)mat. This is a known translocation and is not unique to the proband in this pedigree (39) so we therefore conclude this is not related to the CNS malformations. Whole exome sequencing (Illumina Hi Seq 2000) was performed for the affected child and parents after obtaining informed consent according to our Cincinnati Children’s Hospital Medical Center (CCHMC) IRB-approved protocol. Alignment and variant detection was performed using the Broad Institute’s web-based Genome Analysis Toolkit (GATK; Genome Reference Consortium Build 37) (40). Quality control and data filtering were performed on VCF files independently using manual curation and Golden Helix’s SNP and Variation Suite. The initial bioinformatics analysis identified 144,624 variants (Table 1). After filtering for quality control, coding variants, and minor allele frequencies (variants present in dbSNP) we tested three inheritance models. No compound heterozygous mutations were found and only one recessive variant was identified. The recessive variant is in a gene with known roles in dentin biology (*DSPP*), so we did not consider it causal in this family. Analysis for *de novo* variants identified 7 candidates. Two were not supported by a new analysis of the bam files and considered technical artifacts and another was seen 14 times in an internal cohort and was thus considered a geographically localized variant or a local technical artifact. Three more variants were in genes with known mutations in mouse or human which were not consistent with the proband (*LRGUK, NLRP1, LILRB3*). The final variant was a missense variant in *TUBA1A:* c.1225G>T (p.V409F). This variant was confirmed by Sanger sequencing and was not present in either parent (Fig. 1D). Furthermore, this variant was not identified in the Genome Aggregation Database (gnomAD)(41, 42) or 1000 genomes project (42). The c.1225 G>T variant was predicted as “disease causing” by MutationTaster and “possibly damaging” by PolyPhen-2 (score: 0.932; sensitivity: 0.80; specificity: 0.94) and results in an amino acid change from a Valine to a Phenylalanine at a conserved residue (V409F, Fig. 1E). The affected residue is predicted to be on the exterior of the α-tubulin monomer, near the adjacent β-monomer (Fig. 1F). Thus, we predict the effect of the variant to be on binding of other molecules such as microtubule associated proteins or motor proteins like kinesin, rather than on tubulin monomer interactions.

**Table 1.**
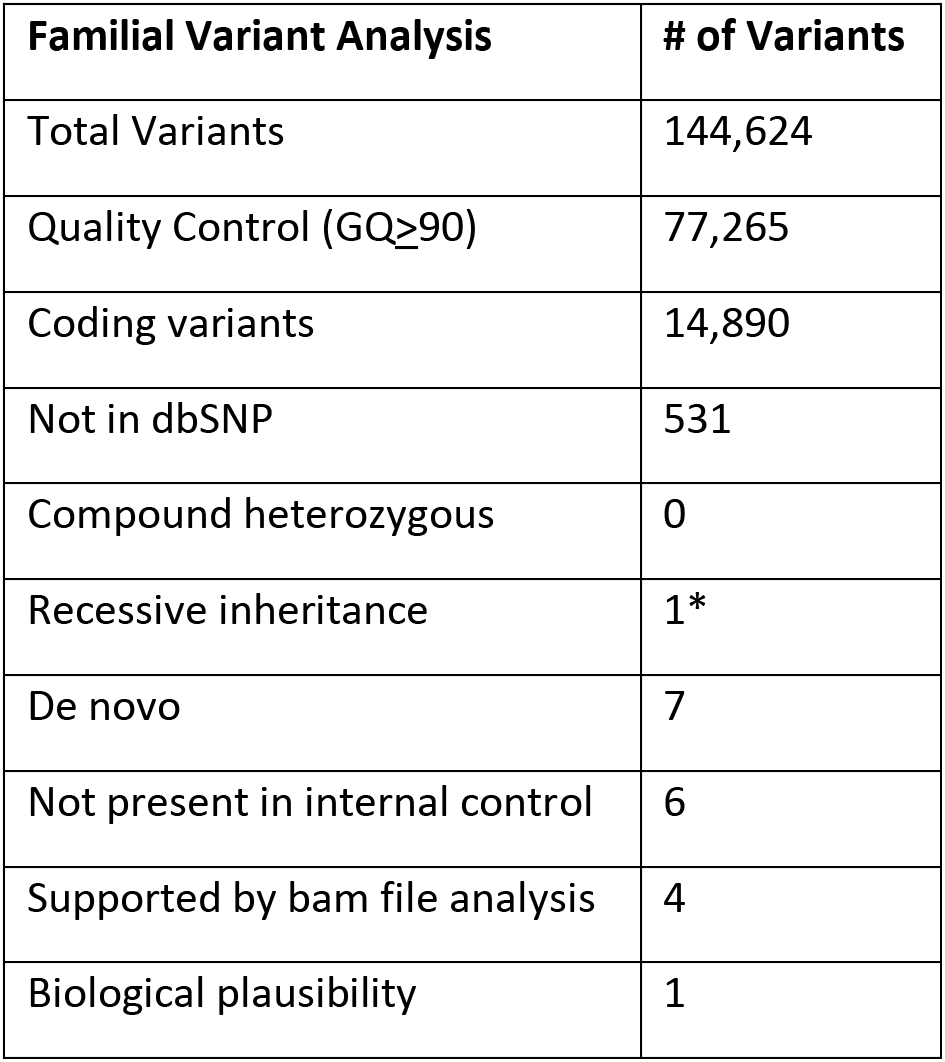
Human Exome Variant Analysis.

While the phenotypes we observed in the proband are within the broad tubulinopathy spectrum, they are among the most severe reported to date for a tubulin mutation (5, 18). *TUBA1A* has been previously associated with neuronal migration defects and lissencephaly; however, these phenotypes were not observed in our patient. Recent findings suggest the spectrum of *TUBA1A*-related brain malformations may be broader than initially thought (18). Shared features include an absence of a septum pellucidum, poor formation of basal ganglia, a significantly small/dysmorphic brain stem, and evidence of hydranencephaly (18). The sheer extent of cortical tissue loss is unique, as is the aqueductal stenosis.

Recent studies have identified other patients with missense mutations in *TUBA1A* at the identical amino acid residue (p.V409A and p.V409I, 2). In the patient with the p.V409I variant, the major finding was only central pachygyria. In contrast, both the p.V409A (2) and p.V409F (this study) patients show severe loss of cortical tissue formation, with the proband in this study more dramatically affected. Of these three missense mutations, the p.V409F variant is the most structurally dissimilar with the introduction of a large phenylalanine aromatic side-chain. Given the extreme phenotypic consequences of this mutation, we suspect this disrupts more of the functions of the tubulin monomer.

### Null alleles of *Tuba1a, Tubb2a, Tubb2b*, and *Tuba8*

The finding in the patient described here continues a pattern in the human genetics of the tubulinopathies wherein pathogenic mutations are found as *de novo* mutations in the affected individuals, and not as rare alleles in the general population. The mechanism of these is unknown but is likely to be a constellation of dominant negative effects. The consequences of loss of function for these molecules has not been tested in a whole animal model. We therefore used CRISPR-Cas9 genome editing to create deletions of *Tuba1a, Tubb2a*, and *Tubb2b*. Guide RNAs were generated to target the endonuclease to two intergenic regions flanking each tubulin gene (Supplemental Table 1). *Tubb2a* and *Tubb2b* are only 49kb apart (0.02cM) in the mouse genome, suggesting that meiotic recombination between the loci will be an extremely rare event. With this consideration in mind, we first attempted to multiplex CRISPR-Cas9 editing for *Tubb2a* and *Tubb2b* with the goal of creating a single deletion of each gene as well as a simultaneous deletion of both. A total of four guides were created for each gene, two for each side of each gene (Fig. 2A,D). A mix of all eight *Tubb2a* and *Tubb2b* guides were constructed and used for blastocyst microinjection. The resulting founders were analyzed by PCR and Sanger sequencing for alterations at the *Tubb2a* and/or *Tubb2b* loci. We recovered two alleles of *Tubb2a* and one of *Tubb2b. Tubb2a^em1Rstot^* (hereafter referred to as *Tubb2a^d3964^*) is a 3,964bp deletion (chr13: 34,074,221 - 34,078,184; GRCh37) and *Tubb2a^em2Rstot^* (*Tubb2a^d4223^*) is a 4,223bp deletion (chr13: 34,074,224 - 34,078,447). Both independent deletions excise the entire *Tubb2a* open reading frame (Fig. 2A-C, S1A). We also obtained one complete deletion allele of *Tubb2b. Tubb2b^em1Rstot^* (*Tubb2b^d4186^*) is a 4,186bp deletion (chr13: 34,126,376 - 34,130,558) (Fig. 2D-F).

**Figure 2.**
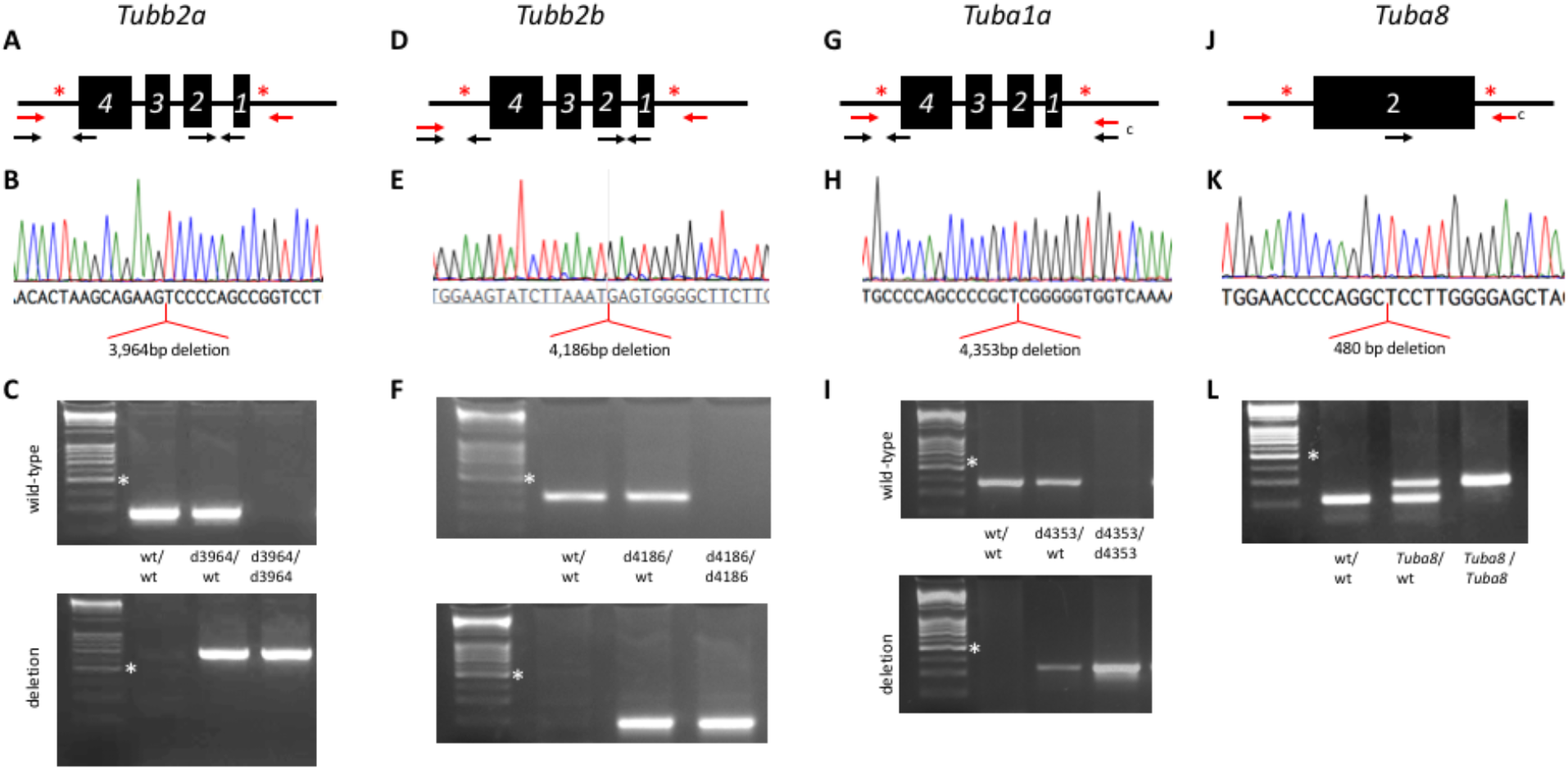
Novel CRISPR/CAS9 alleles of four tubulin genes. Deletion of *Tubb2a* (A-C), *Tubb2b* (D-F), *Tuba1a* (G-I) and exon 2 of *Tuba8* (J-L). (A,D,G,J) Schematics for deletions showing sites of CRISPR guide sequences (red asterisk), arrows indicate relative locations of PCR primers (red are meant to only amplify under standard conditions when gene deletion occurs). (B,E,H,K) Sequencing chromatograms showing the results of mutant specific PCR products. Alignments to the reference genome indicate the deletions indicated for each gene. (C,F,I,L) PCR genotyping to specifically identify each allele. Note *Tubb2a, Tubb2b*, and *Tuba1a* are on the reverse strand of their respective chromosomes. White asterisk indicates 500 bp marker in DNA ladder.

In a parallel manner, four CRISPR guides were generated to delete *Tuba1a* (Fig.2G) and we recovered two independent deletions (Fig.2G-I; S1B,D). *Tuba1a^em1Rstot^* (*Tuba1a^d4353^*) is a 4,353 bp deletion (chr15:98,949,678 - 98,954,031) and *Tuba1a^em2Rstot^* (*Tuba1a^d4262^*) is a 4,262 bp deletion (chr15: 98,949,770 - 98,954,031). The *Tuba8^em1J^* (*Tuba8^delex2^*) allele was obtained commercially and is a CRISPR-Cas9 mediated deletion of exon 2 (Fig. 2 J-L).

### *Tubb2a, Tubb2b*, and *Tuba8* are not required for normal brain morphogenesis

Mice heterozygous for all the deletion alleles were viable and fertile. As we hypothesized the deletion alleles would yield recessive phenotypes, we intercrossed heterozygous carriers for each. We found no reduction from Mendelian expectations in the number of live animals at weaning for any allele of *Tubb2a, Tubb2b*, or *Tuba8* (Table 2) and animals homozygous for these deletions did not appear grossly different than littermates. We collected brains for both whole mount analysis and histological examination from postnatal day (P)28-31 animals to determine if loss of these tubulin genes caused any discernable malformations of cortical development. Whole mount imaging of the brains did not reveal any gross phenotypes (Fig. 3). Histological analysis did not indicate any more subtle anomalies (Fig. 3). During the course of our analysis, a conditional deletion *Tuba8* allele was reported by another group to have no brain phenotype (43). Consistent with this finding, the *Tuba8^delex2/delex2^* homozygous mice we analyzed are viable and fertile. Their brains were also collected from P28-P31 and had no distinct gross morphological changes. H&E and Nissl staining also did not show any morphological differences when compared to wild type animals and we did not explore these any further (Fig. 3 M-O). Body weight at P28 did not show any evidence of a failure to thrive upon loss of *Tubb2a* or *Tubb2b* (Fig. 3P-R). We conclude from these data that neither *Tubb2a, Tubb2b* nor *Tuba8* are uniquely required for cortical developmental or survival in the mouse.

**Figure 3.**
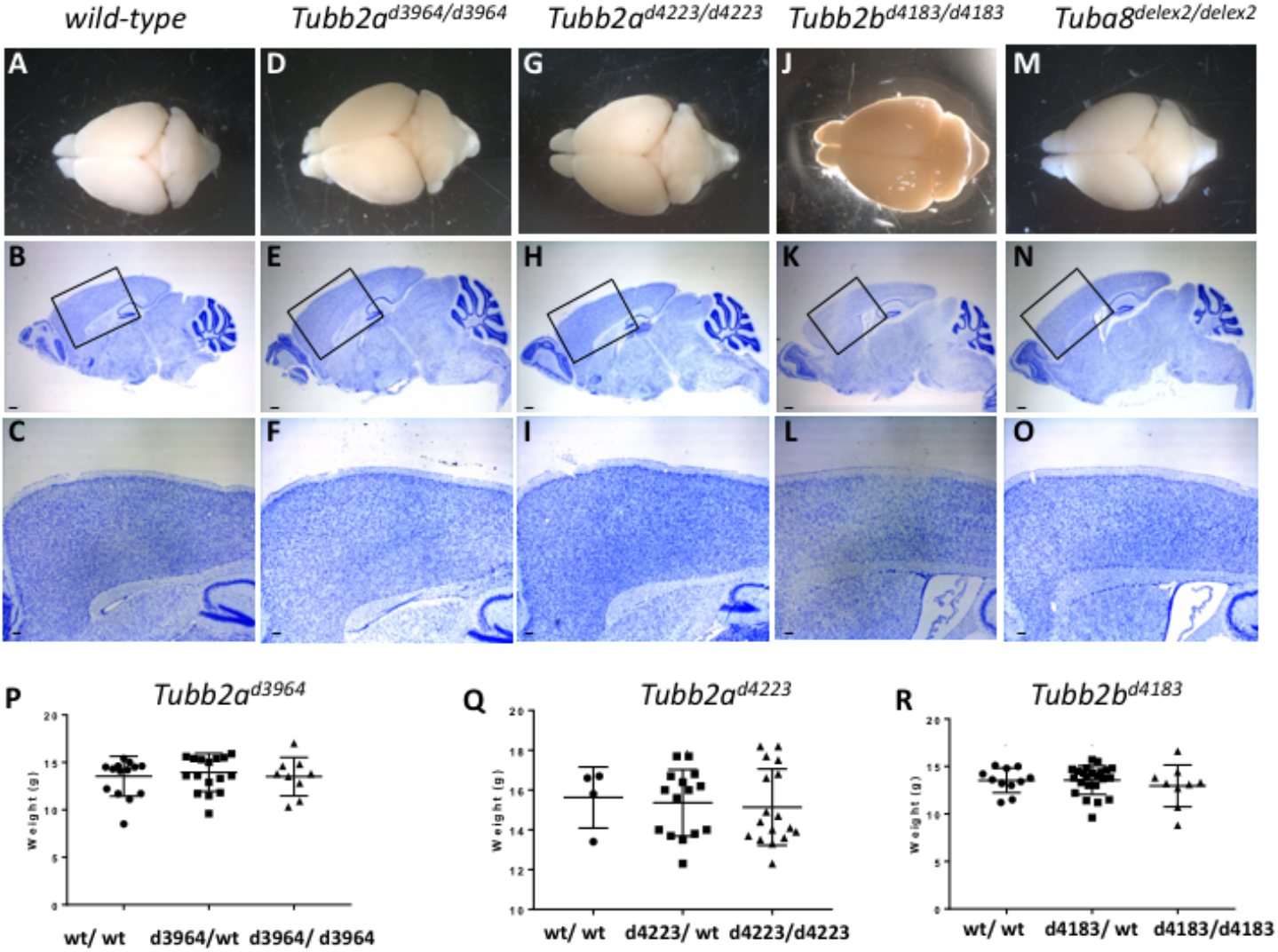
Deletions of *Tubb2a, Tubb2b* or *Tuba8* do not result in lethality, gross brain malformations or failure to thrive. Whole mount and Nissl staining histological analysis of *Tubb2a^d3964/d3964^* (D,E,F), *Tubb2b^d4223/d4223^*(G,H,I), *Tubb2b^d4186/d4186^*(J,K,L), and *Tuba8^delex2/delex2^* (M,N,O) homozygous mice at P28-31. (P-R) Postnatal weights of the *Tubb2a* and *Tubb2b* alleles at P28 show no sign of failure to thrive. Scale bars indicate 100 mm in B,E,H,K,N and 0.2 mm in C,F,I,L,O. Boxes in B,E,H,K,N show regions highlighted in C,F,I,L,O, respectively.

**Table 2.**
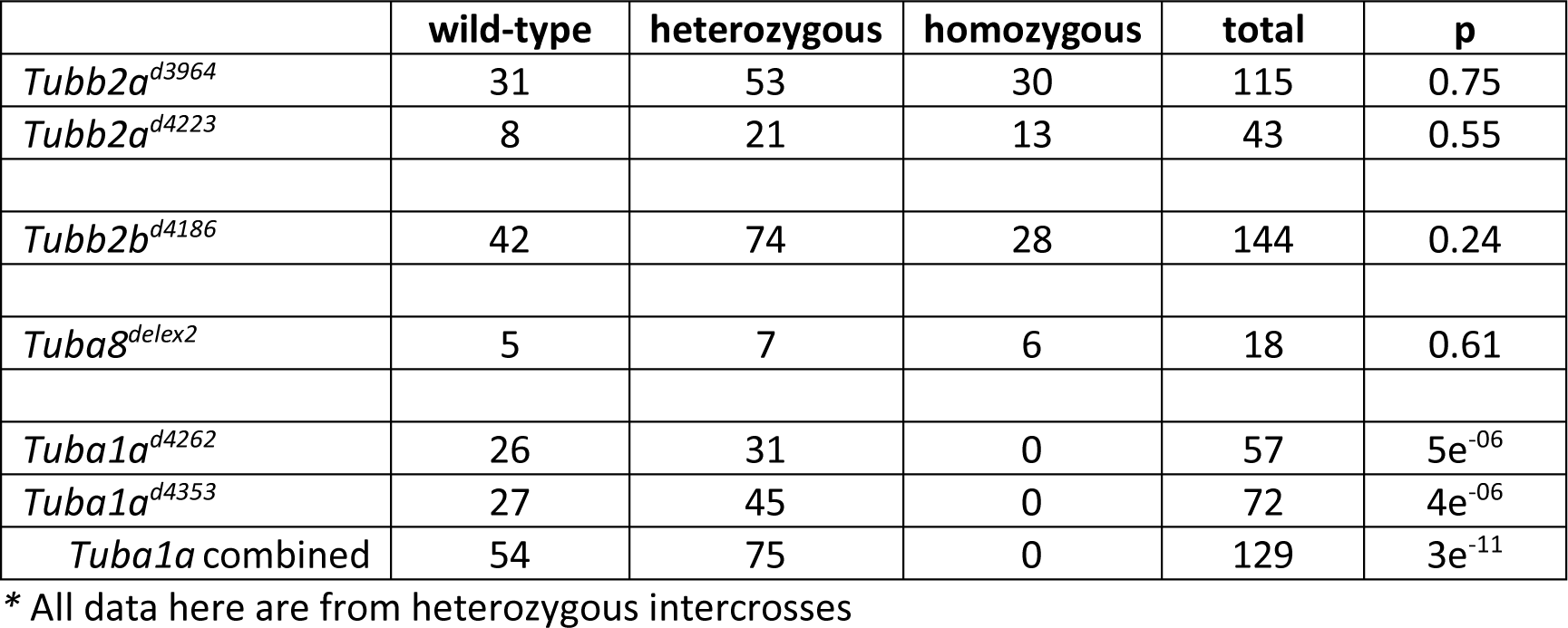
Survival at weaning of mice with deletions of *Tubb2a, Tubb2b, Tuba8* and *Tuba1a\**.

### *Tuba1a* is required for survival and brain development

In stark contrast to our findings with *Tubb2a, Tubb2b*, and *Tuba8, Tuba1a* is absolutely required for survival to weaning as we did not recover any homozygotes for either *Tuba1a* deletion allele (n=129 total; Table 2). We do note a small reduction (approx. 12%) in heterozygote survival among the surviving animals. To begin to understand the reason for the lethality and to assess brain development in the homozygotes, we collected embryos at late organogenesis stages. A combined analysis of both *Tuba1a* deletion alleles in embryos from E14.5-E18.5 did not reveal a significant loss of homozygous embryos during embryonic stages (Table 3).

**Table 3.** Embryonic survival of *Tuba1a* deletion mutants (E14.5-E18.5).

|  | wild-type | heterozygous | homozygous | total | p |
| --- | --- | --- | --- | --- | --- |
| *Tuba1a*<sup>d4262</sup> x *Tuba1a*<sup>d4262</sup> | 7 | 18 | 11 | 36 | 0.64 |
| *Tuba1a*<sup>d4353</sup> x *Tuba1a*<sup>d4353</sup> | 22 | 68 | 27 | 117 | 0.17 |
| *Tuba1a* combined | 29 | 86 | 38 | 153 | 0.18 |

\* All data here are from heterozygous intercrosses

Whole mount images of *Tuba1a^d4353/d4353^* and *Tuba1a^d4262/d4262^* animals had very obvious and consistent phenotypes. We noticed frequent thoracic edema and significant hemorrhaging in the cervical region extending towards the forelimbs (Fig. 4A,F,K). Histological analysis of the developing forebrain revealed a striking series of cortical malformation phenotypes. Sections from both posterior and anterior regions of the forebrain highlight enlarged cerebral and third ventricles, as well as a widened base of the third ventricle in posterior regions. The basal ganglia are reduced in size in the mutants as compared to control. Examination of cortical morphology showed a reduction in the cortical plate and a possible increase in the width of the ventricular zone at E16.5. The width of the intermediate zone also appears decreased. (e.g. Fig. 4M). We also noted a number of the homozygous mutants had cleft palate (Fig. 4Q) This was incompletely penetrant and seen in 3/11 *Tuba1a^d4353/d4353^* embryos (27%) and 2/16 *Tuba1a^d4262/d4262^* (12.5%) embryos. We noted cleft palate in only one out of 52 heterozygous embryos analyzed, but we suspect this is likely due to delayed development in that particular embryo based on morphological staging. We conclude that *Tuba1a* is required for survival and that loss of *Tuba1a* leads to major cortical malformations with similarities to what is seen in the human tubulinopathy patients.

**Figure 4.**
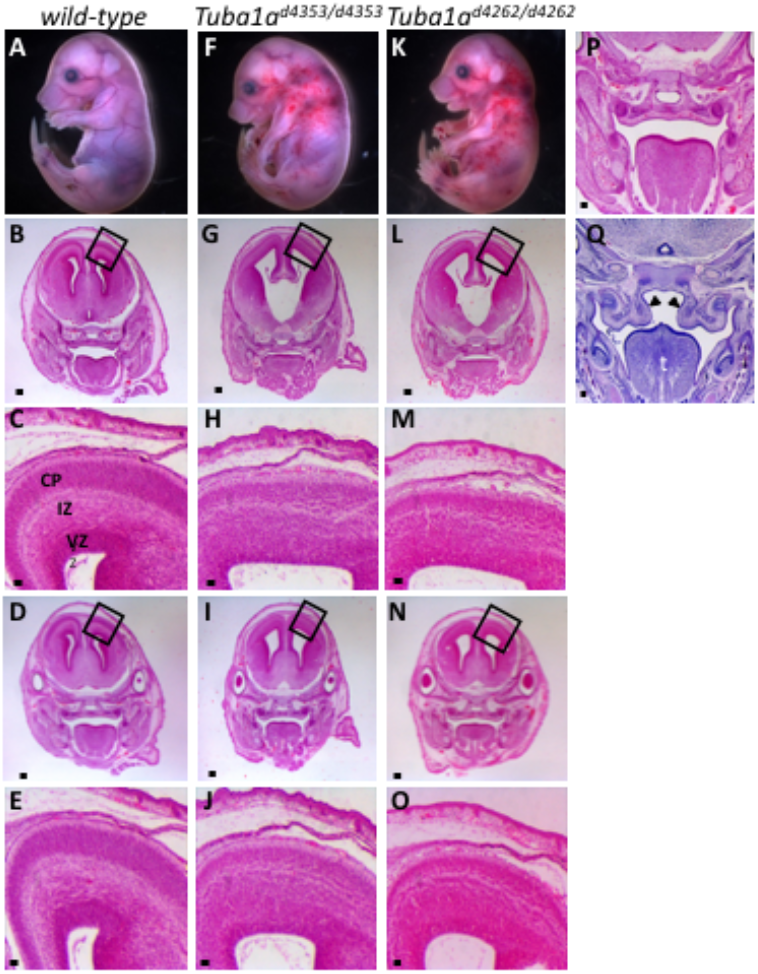
Loss of *Tuba1a* leads to perinatal lethality and cortical malformations. Embryos homozygous for either of two *Tuba1a* deletions (*Tuba1a^d4353^* F-J; *Tuba1a^d4262^* K-O) have significant gross morphological phenotypes when compared to wild-type (A-E). Cervical hemorraghing is often noted (F,K). Hypoplastic basal ganglia and disruptions to the cortical plate (CP), intermediate zone (IZ) and ventricular zone (VZ) are noted in both mutants. making them distinguishable upon initial embryo harvesting. Histological analysis shows enlarged ventricles with loss of IZ. Black boxes in B,G,L show ighlighted areas in C,H,M, respectively. Some mutants show cleft palate (Q) resulting from a failure of the palatal shelves (arrowheads) to elevate and fuse as seen in wild-type embryos (P). Scale bars indicate 200 μm in A, G,L,D,I,N; 50 μm in C,H,M,E,J,O; and 100 μm in P,Q.

### A novel ENU allele of *Tuba1a*

A separate experiment utilizing an ENU mutagenesis forward genetic screen to identify new alleles important for cortical development identified an additional mutant with similar brain phenotypes. The *quasimodo* mutants were first identified by an abnormal curvature of the thoracic region seen in many of these mutants (Fig. 5D). Histological analysis of mutants showed a significant brain malformation with cortical ventriculomegaly, a reduced ventricular zone and cortical plate accompanied by a reduced intermediate zone and enlarged third ventricle (Fig. 5E,F). Whole exome sequencing of three homozygous mutants identified a large number of homozygous variants as predicted. Filtering for SNPs which were shared by all three mutants, not present in dbSNP, predicted to have a high/moderate effect on the protein, and having only one variant in the gene (i.e., not a highly polymorphic sequence) left a total of nine variants. A review of literature to identify known consequences for loss of function in these genes or known roles of these genes did not yield any compelling candidates (Table 4).

**Figure 5.**
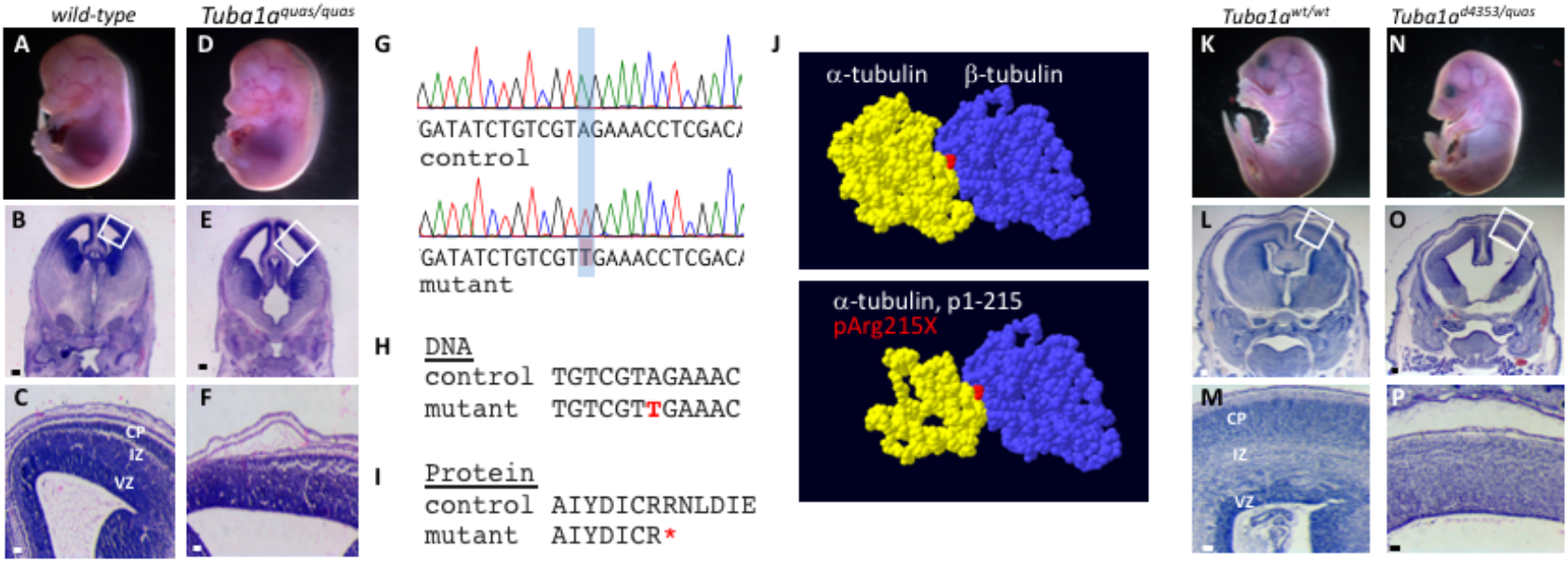
The *quasimodo* mutation is an allele of *Tuba1a*. (A-F) Homozygous *quasimodo* mutants are distinguishable at E14.5 by the systemic edema (D). Histological analysis show enlargement of the third ventricle (E) and reduced cortical tissue (F). (G) Sanger sequencing confirms homozygosity for a candidate SNP in a conserved DNA (H) and protein (I) sequence (SNP and coding change shown in red, red asterisk indicates premature stop codon). (J) Structure of the wild-type α/β tubulin dimer is shown above and the remaining α-tubulin structure is shown below (red = non-hydrolyzable GTP at the monomer interface). (K-P) Complementation analysis with the *Tuba1a^d4353^* is consistent with *quasimodo* being an allele of *Tuba1a* as *Tuba1a^d4353/quas^* mutants have gross (N) and histological phenotypes similar to the other *Tuba1a* homozygous phenotypes. The failure of alleles to complement indicates both mutations are in Tuba1a Boxes in B,E,L,O show areas enlarged in C,F,M,P, respectively. Scale bars indicate 200 μm in B,E,L,D and 50 μm in C,F,M,P.

**Table 4.** Mouse Exome Variant Analysis.

| Exome Variants | # of variants |
| --- | --- |
| total | 199085 |
| Homozygous in all three animals | 982 |
| not in dbSNP | 575 |
| SNP | 324 |
| High/ moderate predicted effect | 43 |
| Single variant in gene | 9 |

We then performed a parallel analysis for all heterozygous missense variants shared by the three sequenced mutants with the alternative hypothesis that the *quasimodo* phenotype may be an incompletely penetrant heterozygous variant. One variant from this analysis was a nonsense mutation in *Tuba1a* creating a premature stop codon at amino acid 215 (p.R215*). We noted that the sequence in this region of *Tuba1a, Tuba1b* and *Tuba1c* is highly similar and carefully analyzed the exome .bam files. Virtually all of the reads for this region of genomic sequence were aligned to the *Tuba1a* gene and virtually no reads were mapped to *Tuba1b* or *Tuba1c*. Given the phenotypic similarity to the *Tuba1a* deletion, we hypothesized the similarity among these genes may have confounded the automated sequence alignment and pursued Sanger sequencing with intronic primers that would specifically amplify *Tuba1a*. This analysis showed that the *quasimodo* mutants were all in fact homozygous for the variant creating the nonsense mutation (Fig. 5G-I).

In order to confirm that the *Tuba1a* R215* variant is actually the causative variant in the *quasimodo* mutants, we performed a complementation test with the *Tuba1a^d4353^* deletion allele. We crossed *Tuba1a^quas/+^* with *Tuba1a^d4353/+^* animals and in three litters (22 live pups), we recovered no *Tuba1a^d4353/quas^* live animals at weaning (Table 5). We further explored this hypothesis with embryonic dissections and recovered two *Tuba1a^d4353/quas^* embryos at E17.5. These mutants exhibited the exaggerated curvature of the thoracic region, as previously seen in *Tuba1a^quas/quas^* mice (Fig. 5N). Histological analysis showed ventriculomegaly with disrupted cortical architecture including a loss of the intermediate zone, hypoplastic basal ganglia, and cleft palate in both mutants (Fig. 5O,P). The similarity of these phenotypes to the *Tuba1a^d4353/d4353^*and *Tuba1a^quas/quas^* mutants suggests that the R215* variant found in *quasimodo* mutant is indeed the causal variant. Moreover, this would support the above conclusion that loss of *Tuba1a* results in significant cortical malformations. This mutation is predicted to produce a protein missing approximately half of the amino acid sequence, if that protein is even stable in the cellular context (Fig. 5J). The mutant protein would then be missing major portions which would normally interface with the adjacent β-tubulin monomer.

**Table 5.** Quasimodo is an allele of *Tuba1a*.

|  | wild-type | *quas*/wt | d4353/wt | d4353/*quas* | total | p |
| --- | --- | --- | --- | --- | --- | --- |
| *Tuba1a*<sup>d4353/wt</sup> x *Tuba1a*<sup>*quas*/wt</sup><br>weaning | 7 | 6 | 9 | 0 | 22 | 0.04 |
| *Tuba1a*<sup>d4353/wt</sup> x *Tuba1a*<sup>*quas*/wt</sup><br>embryos | 6 | 9 | 8 | 2 | 25 | 0.42 |

## DISCUSSION

We have identified a novel human variant and generated a series of mouse alleles which together significantly add to our understanding of the role of tubulin genes in malformations of cortical development. The human *TUBA1A* p.V409F allele is a variant in a residue previously reported twice in the literature (2) but this specific missense mutation results in a much more severe phenotype. We also identified a new *Tuba1a* mouse allele with severe CNS phenotypes through an ENU mutagenesis experiment. These alleles contribute to a larger body of work indicating the importance of tubulin genes in cortical development and disease. We further created a series of deletion alleles in mouse to determine if *Tuba1a, Tuba8, Tubb2a* and/or *Tubb2b* are essential for development, or if the mutations identified in human genetics are indeed acting as dominant negative heterozygous mutations. We find that *Tuba8, Tubb2a*, and *Tubb2b* can all be deleted from the mouse with no obvious effect on cortical development, animal survival, or health. *Tuba1a*, however, is quite different, as loss of *Tuba1a* is lethal in the mouse and leads to severe malformations of cortical development. Thus, other tubulin genes with high sequence similarity cannot compensate for *Tuba1a* but may be doing so for *Tubb2a* and *Tubb2b*. Together, these data show that some tubulin genes are absolutely critical for CNS development and organism survival while others are dispensable.

*Tubb2a* and *Tubb2b* are highly similar and immediately adjacent in the genomes of both mice and humans, suggesting that a recent duplication event in a common ancestor may have rendered these genes redundant for each other. This may explain why a single deletion of each does not lead to a CNS phenotype. This model can be tested with a double mutant analysis. The genes are sufficiently close to each other within the genome, this experiment will require creation of an independent deletion or conditional allele in *cis* to one of the existing mutations.

*Tuba1a* is highly similar to *Tuba1b* (99.6% identical) and *Tuba1c* (98.2%) and are also clustered in the genome suggesting these may also be able to compensate for each similar to our hypothesis for *Tubb2a* and *Tubb2b*. In direct contrast to our findings with *Tubb2a* and *Tubb2b*, however, loss of *Tuba1a* is catastrophic for the mouse embryo. The individual requirements for *Tuba1b* and *Tuba1c* remain unexplored and we are also addressing this hypothesis with future deletion alleles.

It is not known why the paradigm is so different for these two gene clusters (*Tuba1a/b/c* as compared to *Tubb2a/b*). Do cells express a mix of α and β tubulin monomers? Are there changes in specific gene expression over time, perhaps between neurogenesis stages and terminal differentiation? These questions remain largely unanswered although a *Tubb2b-eGFP* transgenic line suggests *Tubb2* is expressed in both progenitors and postmitotic neurons (44). A detailed expression analysis would be useful in generating models to explain these genetic findings. Unfortunately, this is currently quite challenging. No antibodies will distinguish between these proteins in an immunohistochemical analysis. RNA expression studies are complicated by this homology as well. RNA *in situ* hybridization probes against the untranslated regions may be able to distinguish between genes but some hypotheses about coexpression at the cellular level would be challenging with *in situ* hybridization-based approaches. RNA-Seq, both bulk sequencing and single cell sequencing, would potentially be useful in addressing these models at least in a preliminary way. However, our experience with the *quasimodo* mouse mutant exome sequencing suggests the current library preparations and/or sequence alignment algorithms are confounded by the sequence similarities between these genes and these data sets are to be interpreted with extreme caution. We propose that a series of epitope-fusion protein alleles in the mouse would be a useful tool set to explore both expression of discrete tubulin genes and will serve as a platform for biochemical approaches to identify microtubule associated protein and other cellular interaction partners.

While the phenotypes we present here are striking malformations of cortical development, we do not yet know the cellular and molecular basis for these phenotypes. The histological changes in the ventricular zone, intermediate zone and cortical plate all suggest fundamental perturbations of neurogenesis and/or cellular migration. Detailed mechanistic studies are needed to determine how these processes are altered in the face of tubulin mutations. The human missense mutations are not extensively modeled in mice, with the exception of the *Tubb3* R262C model of CFEOM (38). A mouse model of some of the most common mutations in *Tuba1a* would be helpful in both understanding the underlying mechanism(s) but also as a platform for testing potential therapeutic intervention.

Human genetics studies have identified a series of tubulin genes as crucial building blocks of the brain and tubulins have long been appreciated as important components for neuronal cell biology. As our genomic tools and mouse modeling capabilities have advanced, we have the opportunity to return to some fundamental questions about the requirement(s) and role(s) of tubulin genes in mammalian development and disease.

## Acknowledgements

We are grateful to Yueh-Chiang Hu and the CCHMC Transgenic Animal and Genome Editing Core for adviceand technical assistance, and Bill Dobyns for extensive discussion of this work prior to publication.

## Funding

Cincinnati Children’s Research Foundation and National Institutes of Health (R01NS085023 to R.W.S.)

## METHODS

### Patient sequencing and variant confirmation

Informed consent/assent was obtained from all subjects according to Cincinnati Children’s Hospital Medical Center (CCHMC) institutional review board protocol #2014-3789. All methods and experimental protocol were carried out in accordance with relevant guidelines and regulations and with approval from the CCHMC Institutional Biosafety Committee. Following consent, whole blood was collected on the parents, residual DNA and fibroblasts from the proband. Library generation, exome enrichment, sequencing, alignment and variant detection were performed in the CCHMC Genetic Variation and Gene Discovery Core Facility (Cincinnati, OH). Briefly, sheared genomic DNA was enriched with NimbleGen EZ Exome V2 kit (Madison, WI). The exome library was sequenced using Illumina’s Hi Seq 2000 (San Diego, CA). Alignment and variant detection was performed using the Broad Institute’s web-based Genome Analysis Tookit (GATK)(40). All analyses were performed using Genome Reference Consortium Build 37.

### Variant Filtering and Pathogenicity Assessment

Quality control and data filtering were performed on VCF files in Golden Helix’s SNP and Variation Suite (Bozeman, MT). Non-synonymous coding variants were compared to three control databases, including NHLBI’s ESP6500 exome data (45), the 1000 genomes project (42), EXAC Browser (46) and an internal CCHMC control cohort (47). Remaining variants were subject to autosomal recessive analysis with emphasis on homozygous recessive variants found in the region of homozygosity identified by SNP microarray. The identified variant was compared to known disease genes in the OMIM and Human Gene Mutation (HGMD) (48) databases, and to reported variants in dbSNP (49) and the Exome Variant Server. The variant was also analyzed using Interactive Biosoftware’s Alamut v2.2 (San Diego, CA) to determine location of mutation within a protein domain, the conservation of the amino acid, the Grantham score (50) and the designation of the mutation by three existing in-silico software tools, SIFT (51), Polyphen (52) and Mutation Taster (53).

### Mouse allele generation

CRISPR guides flanking the tubulin genes of interest were evaluated using the Fusi (Benchling.com) and Moreno-Mateos (Crisprscan.org) algorithms. Potential guide RNA (gRNA) sequences were selected and ordered as complementary oligonucleotide pairs with BbsI over-hangs (Table S1) (IDT, Coralville, IA). These were ligated into the pSpCas9(BB)-2A-GFP (px458) vector and transfected into MK4 cells at low confluence using the Lipofectamine 2000 transfection reagent (Thermo Fisher Scientific, Massachusetts). Cells were harvested 48 h after transfection and genomic DNA was isolated and used with the Surveyor mutation detection kit (IDT) in order to test gRNA cutting efficiency. As a control, cutting efficiencies of potential guides were compared with that of a previously-published mTet2 gRNA. Cas9 and gRNAs were injected into C57BL/6N zygotes (Taconic) by the CCHMC Transgenic Core. Potential founders were validated with Sanger sequencing of tail DNA and subsequently maintained on a C57BL/6J (Jackson Labs) background. pSpCas9(BB)-2 A-GFP (PX458) was a gift from Feng Zhang (Addgene plasmid # 48138). The *Tuba8^em1J^* allele was purchased from Jackson Labs (Bar Harbor, ME).

### Animal husbandry

All animals were housed under an approved protocol in standard conditions. All euthanasia and subsequent embryo or organ harvests were preceded by Isoflurane sedation. Euthanasia was accomplished via dislocation of the cervical vertebrae. For embryo collections, noon of the day of vaginal plug detection was designated as E0.5. Brains were removed and imaged using a Zeiss Discovery.V8 stereo-microscope. Box plots were generated with Prism7 (GraphPad).

### Histological analysis

For adult histology, littermate animals underwent transcardial perfusion using cold heparinized phosphate buffered saline (PBS) and formalin (SIGMA) solution. Brains were dissected and fixed for 72 h in formalin at room temperature followed by immersion in 70% ethanol (for histology). Embryo samples were fixed in Bouin’s fixative solution. Samples were then paraffin embedded, sectioned at 6μm for adult tissue and 10μm for embryonic tissue, and processed through hematoxylin and eosin (H&E) or Nissl staining. Sections were sealed using Cytoseal Mounting Medium (Thermo-Scientific).

All data presented here are available to the community in accordance with best practices for data sharing.

---

**Fig S1.**
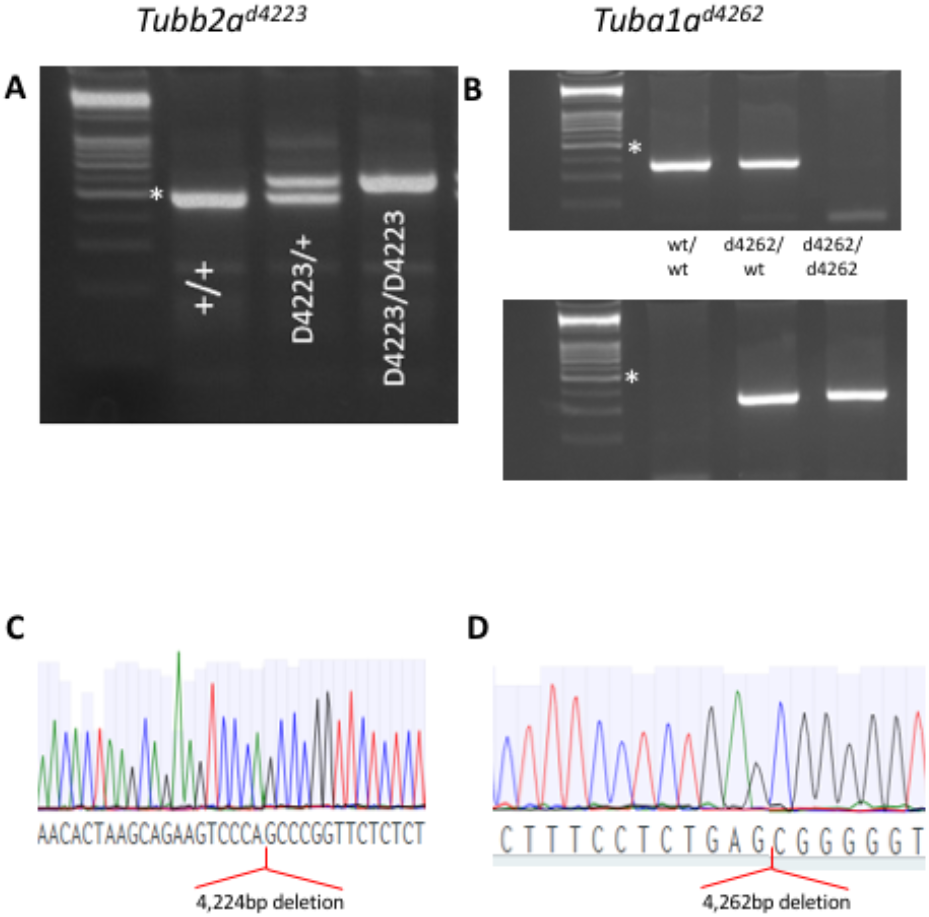
Additional CRISP-Cas9R deletions of *Tuba1a* and *Tubb2a*. PCR genotyping and sequencing results showing deletions sites of two additional alleles of *Tubb2a* (A,C) and *Tuba1a* (B,D).

**Supplemental Table 1.**
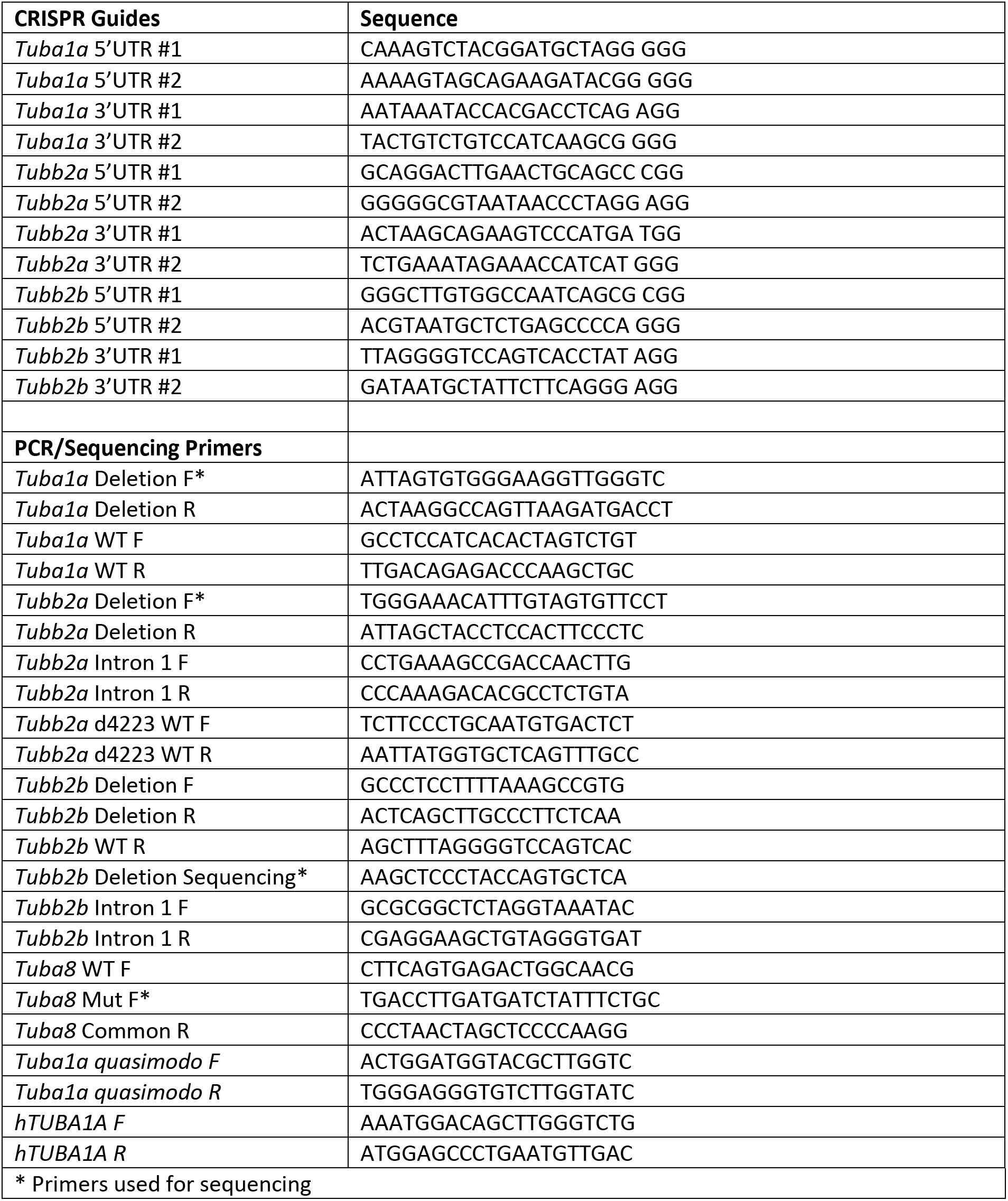
CRISPR guide and PCR primers sequences.

